# When DNA gets in the way in RNA-seq experiments, a sequel

**DOI:** 10.1101/2020.09.28.316265

**Authors:** Jasper Verwilt, Maria D. Giraldez, Wim Trypsteen, Ruben Van Paemel, Katleen De Preter, Pieter Mestdagh, Jo Vandesompele

**Affiliations:** Department of Biomolecular Medicine, Ghent University, 9000 Ghent, Belgium; OncoRNALab, Cancer Research Institute Ghent, 9000 Ghent, Belgium; Digestive Diseases Unit, Virgen del Rocio University Hospital, 41013 Seville, Spain; OncoDigest Group, Institute of Biomedicine of Seville (IBiS), 41013 Seville, Spain

## Abstract

Using a newly developed method dubbed SILVER-Seq—enabling extracellular RNA sequencing (exRNA-seq) directly from a small volume of human serum or plasma— Yan et al. recently reported in Current Biology a potential exRNA biomarker for the early diagnosis of Alzheimer’s disease [1]. After the publication of the initial paper describing the SILVER-Seq method [2], we reported our concern regarding potential DNA contamination in their datasets [3]. Although the authors replied they were able to successfully treat RNA samples with DNase to avoid such contamination, they did not address our observations of the majority of reads without evidence of being derived from RNA, nor documented verified absence of DNA after DNase treatment [4]. To assess whether the newly data generated may suffer from DNA contamination, we downloaded the publicly available sequencing data and evaluated two quality control metrics (i.e., fraction of exonic and splice reads), which were not reported in the paper. We found that both quality metrics were much lower than expected for RNA-seq data (6.28% exonic and 0.478% splice reads), in line with our previous findings on the first SILVER-Seq paper. These observations suggest the data and results presented by Yan et al. are affected by DNA contamination, an issue that may be inherent to the SILVER-Seq technology.

---

RNA sequencing (RNA-seq) has transformed transcriptome characterization in a wide range of biological contexts and is increasingly used to study samples with a low RNA concentration, such as human biofluids. Biofluids contain microRNA and other types of sncRNA, fragments of multiple RNA classes (e.g., mRNA, lncRNA, tRNA, mtRNA) and circular RNA [5]. The presence of a variety of exRNA molecules in the human bloodstream and other biofluids has opened up new avenues for the development of minimally invasive biomarkers for a wide range of diseases. However, the explosion in exRNA research has resulted in a growing field lacking standardized protocols, consensus on data analysis, consistent findings and sufficient experimental detail in many publications, which prevents researchers from critically evaluating the quality of the presented results or reproducing the experiments. Besides, performing exRNA-seq experiments without adequate quality controls may result in several issues, one being sample contamination [6].

RNA-seq contaminants can be either external (originating from a different sample or another species) or, although often overlooked, internal (originating from other molecules from the same sample). Endogenous DNA contamination can be particularly troubling as it can be hard to detect unless specific quality control measurements are performed. RNA-seq experiments suffering from DNA contamination can lead to biased results as it affects proper data quantification and normalization. Due to the low concentration of RNA in human biofluids, DNA contamination can be particularly vexing in exRNA-seq, preventing the reliable detection of potential biomarkers.

DNase treatment is included in most standardized RNA-seq protocols but, in some instances, it is not completely effective (not all DNA is removed) and can result in impaired final libraries. This problem can be aggravated in protocols using crude biofluids without RNA purification, which may contain DNase-inhibiting molecules (one of them being actin, which has long been known for inhibiting DNase activity [7]). Serum in particular contains not only cell-free DNA but also genomic DNA that originates from lysis of white blood cells during *ex vivo* clotting [8], thus increasing the risk of DNA contamination in RNA-seq experiments.

To evaluate whether the RNA-seq signal in the paper by Yan et al. [1] might be affected by contaminating DNA, we replicated the pipeline used in the paper as accurately as possible (no details were reported regarding parameters of sequencing and data analysis) and calculated several quality control metrics. A step-by-step overview of the used tools can be found in the Supplemental Methods section and the full code is uploaded to GitHub (https://github.com/jasperverwilt/exRNA_contamination).

In order to confirm, or refute, our suspicions, we were mainly interested in two data quality metrics: the fraction of exonic reads (5% in case of sequencing pure DNA) and the fraction of splice reads (0% in case of DNA). Considering all samples, we observe exonic fractions ranging from 4.7% to 25.4%, with a median value of 6.28% (Figure 1A); and splice fractions ranging from 0.206% to 1.27%, with a median value of 0.478% (Figure 1B). In addition to the splice and exonic fractions, we checked the strandedness of the data. If SILVER-Seq would employ a stranded library preparation approach (which we do not know for sure, as it is unreported), and the data turns out to be unstranded, contaminating DNA might be at play (since DNA is double stranded, the reads can originate from both strands). With a strandedness of 100% for perfectly stranded data, and 50% for pure DNA, the observed median strandedness was 49.2%, with individual values ranging from 47.9 to 70.4% (Supplemental Figure 1). These results support the hypothesis of DNA contamination.

**Figure 1:**
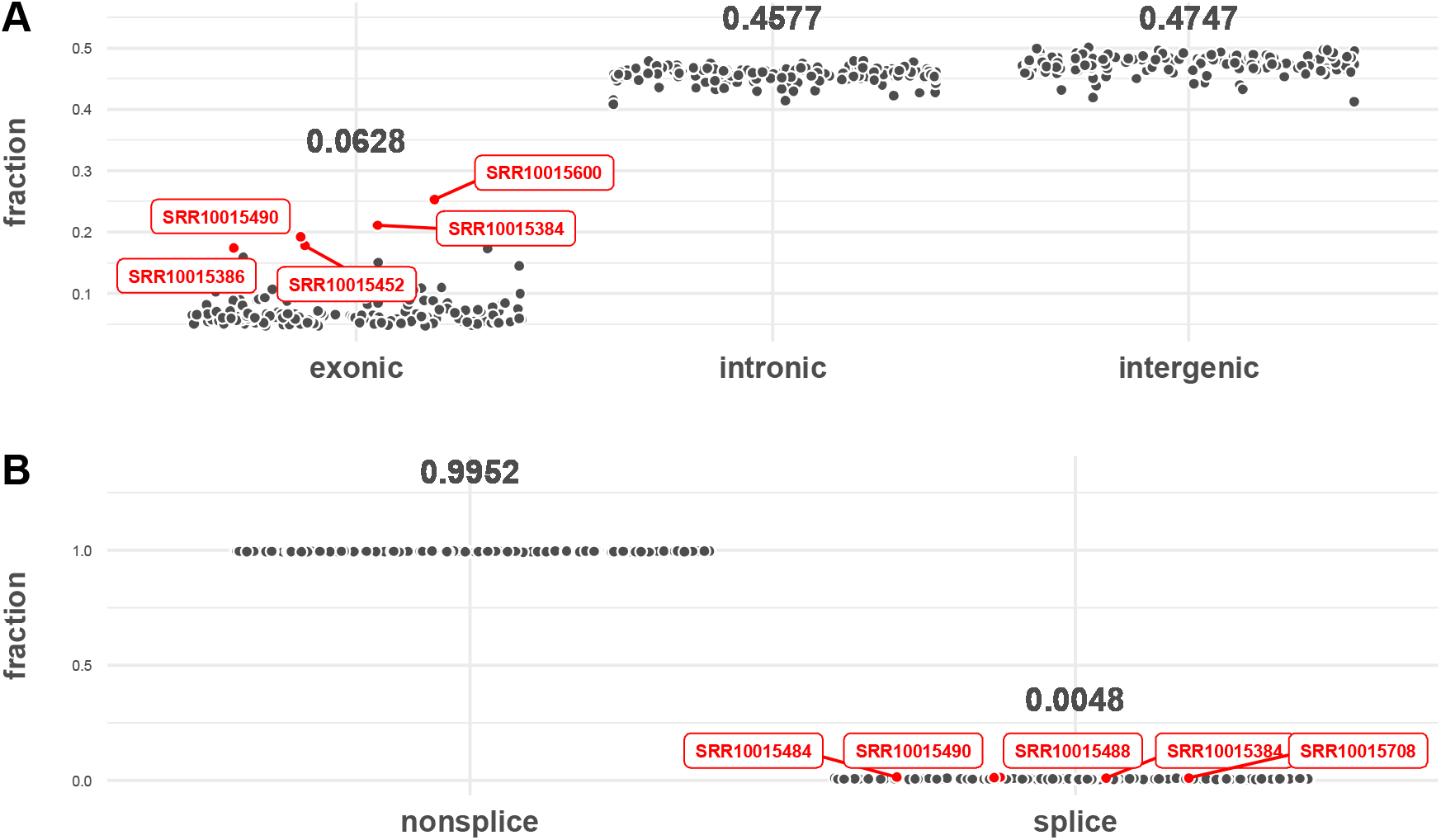
Regional coverage and splice read fractions of the data. *(A)* Fractions of reads mapping to exonic, intronic and intergenic regions. The data points are calculated values for individual samples. The median fractions over all samples are printed. The five samples with the highest exonic coverage are annotated and colored red. *(B)* Fractions of reads mapping to splice and nonsplice regions. The data points are calculated values for individual samples. The median fractions over all samples are printed. The five samples with the highest fraction of reads mapping to splice junctions are annotated and colored red.

The low median value of exonic and spliced reads prompts us to conclude that most of the SILVER-Seq data generated by Yan et al. is affected by DNA contamination. We deduct that SILVER-Seq is a stranded library prep method, given that some samples showed a strandedness higher than 50%. The wide ranges of the exonic and splice read fractions and variable strandedness level indicate that DNA is differentially present in the samples, with some samples performing consistently worse or better for all quality metrics: SRR10015490, for example, showed relatively high values for all the metrics (Figure 1, Supplemental Figure 1).

Finally, the biogenesis of exRNA is not well established yet and some authors argue that biofluids might be enriched in intron and antisense sequences compared with cellular RNAs [9]. However, we are concerned that DNA contamination is the most likely explanation here as: (a) some exRNA-seq studies have consistently reported a high proportion of exonic reads and adequate strandedness [10]; (b) the inherent challenge of avoiding DNA contamination, especially when working with crude biosamples as input; and, (c) the high variability of the evaluated quality control metrics across the reported samples. We would like to emphasize that our observations do not undermine the potential utility of SILVER-Seq. Our letter is a call for thorough reporting of methodology and analysis details including quality control metrics in exRNA-seq studies. We hope that our plea helps to move the exRNA field forward by promoting consistency among laboratories and increasing experimental transparency and reproducibility.

