## Supplemental Material for "When DNA gets in the way in RNA-seq experiments, a sequel"

### Supplemental Methods

First, we downloaded the available data and merged the FASTQ files containing raw read and unique molecular identifier (UMI). We used UMI-tools [2] to extract the UMIs from the merged FASTQ files, to use for later deduplication in the pipeline. Next, we used Trimmomatic [3] to trim the reads of adaptors and low-quality bases. We mapped the trimmed reads using STAR [4], followed by deduplication with UMI-tools [2]. These steps resulted in a BAM file containing all trimmed, deduplicated, mapped reads of each sample. We used RSeQC [5] to calculate the strandedness of the data and the number of reads mapping uniquely to spliced regions, and SAMtools [6] to determine the reads in the BAM file overlapping with BED files containing genomic locations of exonic, intronic and intergenic regions. Please find the exact code on GitHub (https://github.com/jasperverwilt/exRNA_contamination).

*[a]Department of Biomolecular Medicine, Ghent University, 9000 Ghent, Belgium;*
*[b]OncoRNALab, Cancer Research Institute Ghent, 9000 Ghent, Belgium;*
*[c]Digestive Diseases Unit, Virgen del Rocio University Hospital, 41013 Seville, Spain;*
*[d]OncoDigest Group, Institute of Biomedicineof Seville (IBiS), 41013 Seville, Spain;*

### Supplemental Figure 1

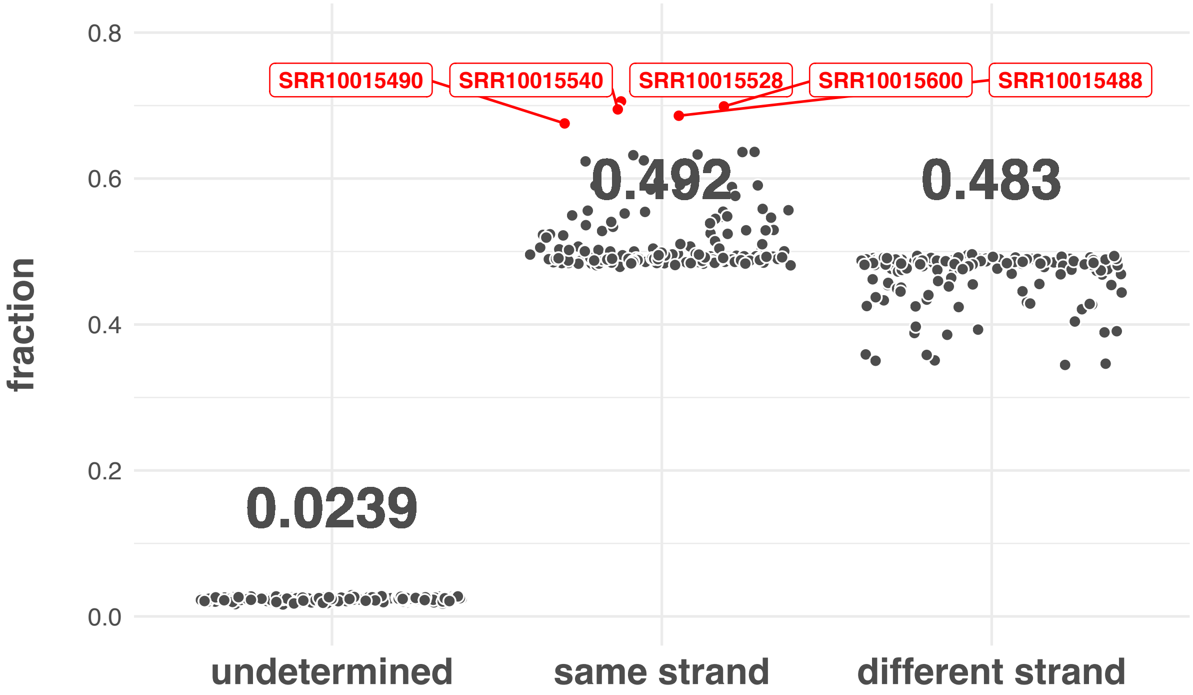


**Supplemental Figure 1:** Strandedness of the data. Reads are categorized according to the strand they map to. ‘Undetermined’ means the read could be transcribed from both ends, ‘same strand’ indicates the reads that mapped to the strand of the annotated parental gene (also named the ‘strandedness’) and ‘different strand’ means the read mapped to the opposite strand of the parental gene. The data points are calculated values for individual samples. The median fractions over all samples are printed. The five samples with the highest strandedness are annotated and colored red.
